# Myeloid DRP1 Sulfenylation Drives Reparative Macrophage Polarization and Neovascularization in Ischemic Muscle

**DOI:** 10.64898/2026.02.15.706043

**Authors:** Shikha Yadav, Sheela Nagarkoti, Varadarajan Sudhahar, Kamarajan Rajagopal, Archita Das, Stephanie Kelley Spears, Tohru Fukai, Masuko Ushio-Fukai

## Abstract

Reparative macrophage polarization and macrophage-derived reactive oxygen species (ROS) are required for ischemia-induced revascularization in peripheral artery disease (PAD). Our previous study showed that mitochondrial fission protein DRP1 promotes reparative polarization and metabolic reprogramming in macrophages and post-ischemic neovascularization. However, the redox-dependent mechanism governing DRP1 activation in this context remains elusive. Here, using a mouse hindlimb ischemia (HLI) model of PAD, we identify cysteine sulfenylation (CysOH) of DRP1 as a critical redox modification induced in ischemic bone marrow (BM)-derived cells. BM chimeric mice reconstituted with CRISPR/Cas9-generated “redox-dead” DRP1-C631A knock-in mutant (*Drp1^C/A^*) BM exhibited markedly reduced limb perfusion recovery and CD31⁺ capillary density in ischemic muscles following HLI. These defects were associated with enhanced Ly6G⁺ neutrophil accumulation, pro-inflammatory F4/80⁺CD80⁺ M1 macrophages and reduced anti-inflammatory F4/80⁺CD206⁺ M2 macrophages in ischemic muscle. Mechanistically, using an in vitro PAD model, hypoxia-serum starvation (HSS) rapidly induced cytosolic ROS production and DRP1-CysOH formation in wild type macrophages. In contrast, *Drp1^C/A^* macrophages failed to undergo DRP1-CysOH-dependent mitochondrial fission under HSS, resulting in aberrant metabolic reprogramming characterized by enhanced glycolysis and mitochondrial ROS, pro-inflammatory p-NF-κB and M1-genes, and suppressed anti-inflammatory p-AMPK and M2-genes. Thus, our findings establish DRP1 sulfenylation as a previously unrecognized redox-sensing mechanism that links ischemia-induced ROS to reparative macrophage reprogramming and revascularization, identifying a novel therapeutic target for PAD.

## 1. Introduction

Peripheral artery disease (PAD) is a common aging-related cardiovascular disorder characterized by impaired ischemia-induced revascularization, chronic inflammation, and progressive tissue damage that can culminate in limb loss.[1]. Despite advances in revascularization procedures, there are no effective therapies that restore endogenous vascular regeneration in patients with critical limb ischemia. A major barrier to therapeutic development is the incomplete understanding of how inflammatory and reparative immune responses are regulated within ischemic tissue [1–4]. Monocyte/macrophage recruitment, polarization, proliferation, and metabolic reprogramming represent critical therapeutic targets for resolving vascular inflammation and enabling revascularization of ischemic muscle [4]. Macrophage phenotypic switching from pro-inflammatory M1-like macrophages that amplify tissue injury and recruit neutrophils, to reparative M2-like macrophages that promote angiogenesis is essential for post-ischemic vascular regeneration [4,5]. The balance between these macrophage states determines whether ischemic tissue undergoes enhanced or impaired regeneration. However, the molecular mechanisms that instruct macrophage fate decisions in response to ischemia remain incompletely defined.

Emerging evidence suggests that reactive oxygen species (ROS) at optimal levels function as signaling molecules rather than merely damaging byproducts and are required for ischemia-induced revascularization. We previously reported that ROS produced from NADPH oxidase 2 (NOX2) in bone marrow (BM) cells and infiltrating immune cells act as key regulators of regenerative myelopoiesis and reparative neovascularization following hindlimb ischemia (HLI), a preclinical model of PAD [6–9]. We also demonstrated that hydrogen peroxide (H₂O₂) produced after growth factor stimulation and HLI promote angiogenesis via cysteine sulfenylation (Cys-OH)-dependent redox signaling [6,10–14]. However, whether and how ROS-mediated redox signaling regulates macrophage reprogramming during ischemia-driven neovascularization remains unknown.

Mitochondria serve as dynamic signaling hubs that govern macrophage oxidative stress, activation, metabolic state, and survival, thereby shaping immune responses. Shifts in mitochondrial oxidative metabolism, mitochondrial ROS (mitoROS) production, tricarboxylic acid (TCA) cycle intermediates, membrane potential, and ultrastructure drive macrophage polarization toward distinct functional states [15]. Pro-angiogenic M2-like macrophage polarization depends on intact mitochondrial respiration, whereas pro-inflammatory M1-like macrophage polarization is associated with enhanced glycolysis [15,16]. The mitochondrial fission GTPase dynamin-related protein 1 (DRP1) is shown to regulate macrophage reprogramming in a context-dependent manner and its activity is regulated by post-translational modifications [17–20]. Using myeloid-specific DRP1 knockout (KO) mice, we previously showed that ischemia-induced Drp1 activation promotes reparative macrophage polarization, metabolic reprogramming, and post-ischemic revascularization [21]. Importantly, ischemia-triggered mitochondrial fission in macrophage occurred independently of canonical DRP1 phosphorylation at Ser616 or Ser637, suggesting non-canonical regulatory mechanism. Because Drp1 contains a redox-sensitive cysteine residue (Cys644 in human; Cys631 in mouse) within the C-terminal GTPase effector domain (GED), we hypothesized that ischemia activates DRP1 in macrophage through a redox-dependent mechanism, thereby promoting reparative macrophage polarization and neovascularization.

In this study using a mouse HLI model of PAD and bone marrow (BM) chimeras reconstituted with CRISPR/Cas9-generated “redox-dead” DRP1^C631A^ knock-in mutant (Drp1^C/A^) mice [14], we identify Drp1 sulfenylation at Cys^631^ in macrophage as a critical redox switch that links ischemia-induced ROS to drive reparative macrophage reprograming and revascularization. Mechanistically, loss of Drp1 sulfenylation in macrophages failed to undergo proper DRP1-dependent fission and metabolic reprogramming characterized by excessive mitoROS production, glycolytic flux, pro-inflammatory p-NfkB and suppressed anti-inflammatory p-AMPK, skewing toward a pro-inflammatory M1-like macrophage phenotype, ultimately impairing reparative neovascularization.

## 2. Materials and Methods

### 2.1. Ethics Statement for Animal Study

All animal studies were carried out following protocols approved by the institutional Animal Care Committee and institutional Biosafety Committee at Augusta University. Room temperature and humidity were maintained at 22.5 °C and between 50% and 60%, respectively. All mice were held under the 12:12 (12-h light: 12-h dark) light/dark cycle. Mice were held in individually ventilated caging with a maximum of 5 or a minimum of 2 mice per cage. A CRISPR/Cas9-engineered “redox-dead” DRP1 knock-in (KI) mouse in which human Cys644 (mouse Cys631) was replaced with alanine (DRP1^C631A^; *Drp1^C/A^*) (Figure 1B) was generated by the transgenic and genomic core facility at Augusta University., as we previously described[14]. Correct targeting of the DRP1 locus was confirmed by PCR and next-generation amplicon sequencing, as reported previously [14]. Homozygous *Drp1^C/A^* KI mice were viable, developed normally, and exhibited body weights comparable to C57BL/6 (wild-type (WT) mice.

**Figure 1:**
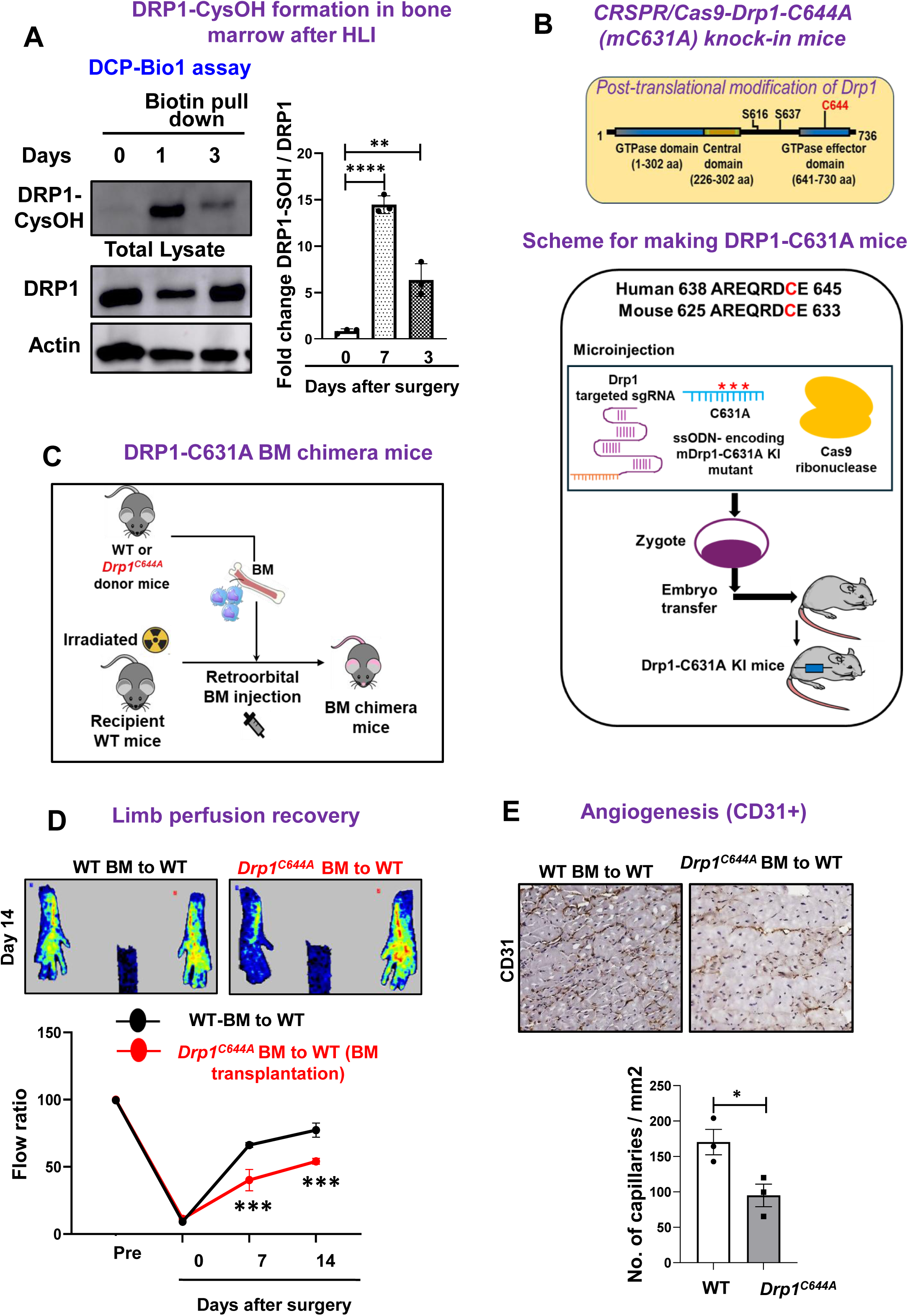
*Drp1^C/A^* bone marrow chimera mice exhibit impaired revascularization after HLI. **(A)** Immunobloting for DRP1Cys-OH in ischemic and non-ischemic bone marrow (BM) of WT mice at indicated days after HLI measured by DCP-Bio1 assay. (**B)** Schematic of CRISPR/CAS9 mediated “redox-dead” mDRP1-C631A (corresponding to human DRP1-Cys644A) knock-in mutant mice generation. **(C)** Schematic of WT and *Drp1^C/A^* bone marrow transplantation (BMT) and hindlimb ischemia. (**D)** Blood flow recovery (ischemic/non-ischemic legs) after HLI measured by laser Doppler flow analyzer. **(E)** CD31^+^ angiogenesis in GC muscle at day 21 after HLI. Data are mean ± SEM (n=3-4) *P<0.05, **P<0.01, ***P<0.001.

### 2.2. Hindlimb ischemia model

Both male and female WT and *Drp1^C/A^* KI mice at 8-15 weeks old were subjected to unilateral hindlimb surgery under anesthesia with intraperitoneal administration of ketamine (87 mg/kg) and xylazine (13 mg/kg). We performed ligation and segmental resection of left femoral artery. Briefly, the left femoral artery was exposed, ligated both proximally and distally using 6-0 silk sutures and the vessels between the ligatures were excised without damaging the femoral nerve. Skin closure was done using 6-0 nylon sutures. We measured ischemic (left)/non-ischemic (right) limb blood flow ratio using a laser Doppler blood flow (LDBF) analyzer (PeriScan PIM 3 System; Perimed) as we described [9,22]. Mice were anesthetized and placed on a heating plate at 37°C for 10 minutes to minimize temperature variation. Before and after surgery, LDBF analysis was performed in the plantar sole. Blood flow was displayed as changes in the laser frequency, represented by different color pixels, and mean LDBF values were expressed as the ratio of ischemic to non-ischemic LDBF.

### 2.3. BM transplantation (BMT)

BMT was performed as published before [22]. Briefly, BM cells were isolated by density gradient separation. Recipient mice were lethally irradiated with 9.5 Gy and received an intravenous injection of 3 million donor bone marrow cells 24 hours after irradiation. Hindlimb ischemia was induced 4-to 6 weeks after bone marrow transplantation.

### 2.4. Histological and Western blot analysis

For cryosections, mice were euthanized and perfused through the left ventricle with saline, limbs were fixed in 4% paraformaldehyde (PFA) overnight and incubated with 30% sucrose, and gastrocnemius muscles were embedded in OCT compound (Sakura Finetek). 7 µm cryosections for capillary density were stained with anti-mouse CD31 antibody (MEC 13.3, BD Bioscience). Macrophages and neutrophils were labeled with anti-F4/80 (BM8, BioLegend) and anti-Ly6G (RB6-8C5, eBioscience) respectively. For immunohistochemistry, we used R.T.U. Vectorstain Elite (Vector Laboratories) followed by DAB visualization (Vector Laboratories). Images were captured by Keyence microscope (bz-x800) or confocal microscopy (Zeiss) and analyzed by Image J or LSM510 software (Zeiss), respectively. Western blot analysis was performed as previously described [6].

### 2.5. Flow cytometry (FACS) analysis

Mouse whole blood and BM cells were collected by cardiac puncture and from femur and tibia respectively using 27G needle in EDTA coated 1mL syringe and placed in 1.5 mL EDTA coated Eppendorf tube for 30 min at room temperature. The peripheral blood was spun down at 500 g for 1 min at 4°C. The supernatant was discarded, and the cells were re-suspended in 1 ml of RBC lysis buffer (00-4300-54, Invitrogen) for 10 min at room temperature in dark. Samples were then washed with DPBS with 2% FCS and centrifuged at 500 g for 5 min at 4°C. Cells were re-suspended in cold DPBS with 2% FCS and placed on ice. Excised gastrocnemius muscles were minced and enzymatically digested in DPBS containing collagenase type I 1 mg/mL Type I (C0130, Sigma), 18 mg/mL collagenase Type XI (C7657, Sigma), 1 mg/mL hyaluronidase (H3506, Sigma) and 50 U/mL DNAse (D4263, Sigma) at 37°C for 1 hour. Cells were spun and re-susupended in DPBS with 2% FCS followed by passing through 70 µm and 40 µm filter and counted. The cells suspension was incubated with blocking buffer consisting of anti-mouse CD16/CD32 (14-0161-85, eBioscince) and 2% FCS for 15 min in ice. Followed by staining with specific antibodies for CD45, CD11b, Ly6G, Ly6C, F4/80, CD206, CD80 (antibody catalog number are mentioned in supplementary table S2) and fixable viability dye (65-0866-14, Invitrogen) at 4°C for 30 min. Specific isotype controls were used to determine gating. The cells were washed and fixed with 4% PFA and acquired on a ThermoFisher Attune Nxt flow cytometer. The recorded data were analyzed using FLowJO 10 software.

### 2.6. Isolation and primary culture of BM- derived macrophages (BMDMs)

BMDMs were harvested from hind leg tibiae and femur of mice. Briefly, BM cells were flushed from the bone with a 27G needle in DPBS and filtered using 70 µm filter. Cells were cultured and differentiated in DMEM medium (Gibco) supplemented with antibiotics, 10% FBS and 20% conditioned media from L929 cell line (enriched in CSF-1) for 7 days in polystyrene culture plates and non-adherent cells were washed and removed every alternate day. The resulting BMDM population was determined by staining with anti-CD11b anti-Ly6C and anti-F4/80 (antibody catalog number are mentioned in supplementary Table S2) and assessed by flow cytometry.

### 2.7. *In Vitro* Hypoxia Serum Starvation (HSS)

Day 7 BMDMs were washed twice to remove traces of FCS and then incubated in starvation medium from Cell applications Inc (Cat: 209-250) and subjected to hypoxia (2% O_2_) for indicated times [23,24].

### 2.8. Quantitative RT-PCR

Total RNA was prepared from cells or tissues using Tri Reagent (Molecular Research Center Inc.). and phenol/ chloroform. Reverse transcription was carried out using high-capacity cDNA reverse transcription kit (Applied biosystems) with 2µg of total RNA. The PCR was performed as per manufacturer’s protocol using ABI Prism 7000 Sequence Detection System 26 (Applied Biosystems, CA) and the QuantiFast SYBR Green PCR kit (Qiagen) for specific genes. Primer sequences for q RT-PCR are enlisted in Supplementary Table 1. The expression of genes was normalized and expressed as fold-changes relative to HPRT.

### 2.9. DCP-Bio1 assay to detect Cys-OH formed (sulfenylated) proteins

To measure sulfenic acid (CysOH) formation (sulfenylation) of proteins, tissues or cells were lysed in degassed-specific lysis buffer [50 mM HEPES, pH7.0 at room temperature, 5 mM EDTA, 50 mM NaCl, 50 mM NaF, 1 mM Na_3_VO_4_, 10 mM sodium pyrophosphate, 5 mM Iodoacetamide (IAA), 100 µM DTPA, 1% Triton-X-100, protease inhibitor, 200 unit/mL catalase (Calbiochem), 200 µM DCP-Bio1 (KaraFast, USA)] and then DCP-Bio1-bound proteins were pulled down with streptavidin beads (Thermo scientific, USA) overnight at 4 °C. DCP-Bio1-bound CysOH formed, sulfenylated-proteins were determined by immunoblotting with specific antibodies, as reported [11].

### 2.10. Statistical analysis

Each experiment was repeated at least 3 times and data are presented as mean ± SEM. Comparison between two groups were analyzed by unpaired two tailed Student *t*-test. Experiments with more than 2 subgroups were analyzed by ANOVA followed by the Tukey post-hoc or Bonferroni multiple comparison analysis to specify the significance between group differences. Values of *p<0.05, **p<0.01, ***p<0.001 were considered statically significant. Statistical tests were performed using Graphpad Prism v10 (GraphPad Software, San Diego, CA).

## 3. Results

### 3.1 Myeloid DRP1 sulfenylation is required for ischemia-induced reparative angiogenesis in response to HLI

We previously demonstrated that myeloid DRP1 and BM-derived ROS are essential for post-ischemic reparative angiogenesis in the HLI model of PAD [6,7,13,21].. Notably, ischemia activates DRP1-dependent mitochondrial fission in macrophages without increasing canonical DRP1 phosphorylation, suggesting a non-phosphorylation-based activation mechanism. We therefore investigated whether ischemia induces cysteine oxidation of DRP1 *in vivo* and whether this modification is functionally required for reparative neovascularization. Using the HLI model, DCP-Bio1 assay revealed that DRP1 sulfenylation (DRP1-CysOH) in BM cells was markedly increased at day 1 post-HLI compared with non-ischemic BM cells, which was gradually declined by day 3 (Figure 1A). To define the functional significance of endogenous DRP1 cysteine oxidation, we generated a CRISPR/Cas9-engineered “redox-dead” Drp1^C631A^ (*Drp1^C/A^*) KI mice (Figure 1B), as described[14]. To specifically assess the role of myeloid DRP1 sulfenylation in post-ischemic angiogenesis, we performed BM transplantation (BMT) to reconstitute lethally irradiated WT recipients with BM from either WT or *Drp1^C/A^* KI donors (Figure 1C). Laser speckle contrast imaging revealed that perfusion recovery at days 7 and 14 following HLI was significantly impaired in *Drp1^C/A^*-to-WT BM chimeric mice compared with WT-to-WT controls (Figure 1D). Consistent with these findings, immunohistochemical analysis of ischemic muscle demonstrated a marked reduction in CD31-positive capillary density in *Drp1^C/A^* -to-WT mice after HLI (Figure 1E). Collectively, these results demonstrate that myeloid DRP1 sulfenylation is required for effective perfusion recovery and reparative neovascularization following ischemic injury, identifying DRP1 cysteine oxidation as a critical redox-dependent mechanism linking ischemia to macrophage-driven angiogenesis.

### 3.2. Myeloid DRP1 sulfenylation promotes pro-angiogenic M2-like macrophage polarization in ischemic muscle

To determine whether myeloid DRP1 sulfenylation regulates immune cell infiltration, macrophage differentiation, and polarization during ischemic injury, we performed flow cytometry-based immunophenotyping of ischemic GC muscles from WT and *Drp1^C/A^* KI BM chimera mice at days 3 and 7 following HLI (Figure 2A). Immunofluorescence analysis revealed a decrease in total number of F4/80^+^ macrophages in GC muscle of *Drp1^C/A^* KI BM chimera than the WT mice (Figure 2B). Further, flow cytometry analysis confirmed that the number of F4/80^+^CD64^+^ mature macrophages was significantly reduced at day 3, but not at day 7, after HLI in *Drp1^C/A^* KI BM chimeric mice compared with WT controls (Figure 2C-D). Notably, inhibition of myeloid DRP1 cysteine oxidation resulted in a marked shift in macrophage polarization. The number of F4/80^+^CD64^+^CD206^+^ M2-like pro-angiogenic macrophages significantly decreased, whereas F4/80^+^CD64^+^CD80^+^ M1-like pro-inflammatory macrophages significantly increased at both days 3 and 7 after HLI in ischemic muscle of *Drp1^C/A^* KI BM chimera mice relative to WT mice (Figure 2E-F). We next assessed whether myeloid DRP1-CysOH reduction alters monocyte or neutrophil recruitment to ischemic tissue. While Ly6G^+^ neutrophils were significantly increased at day 3 after HLI in *Drp1^C/A^*KI BM chimera mice compared with WT mice, their numbers declined by day 7 (Figure S1A-B). In contrast, Ly6Chi monocyte numbers were comparable between genotypes at both days 3 and 7 following HLI (Figure S1A-B). Together, these findings indicate that myeloid DRP1 sulfenylation is dispensable for monocyte recruitment but is required for proper macrophage polarization toward a pro-angiogenic, anti-inflammatory M2-like phenotype in ischemic muscle. Loss of DRP1-CysOH skews macrophages toward an M1-like phenotype, providing a mechanistic link between impaired macrophage reprogramming and defective ischemia-induced revascularization after HLI.

**Figure 2:**
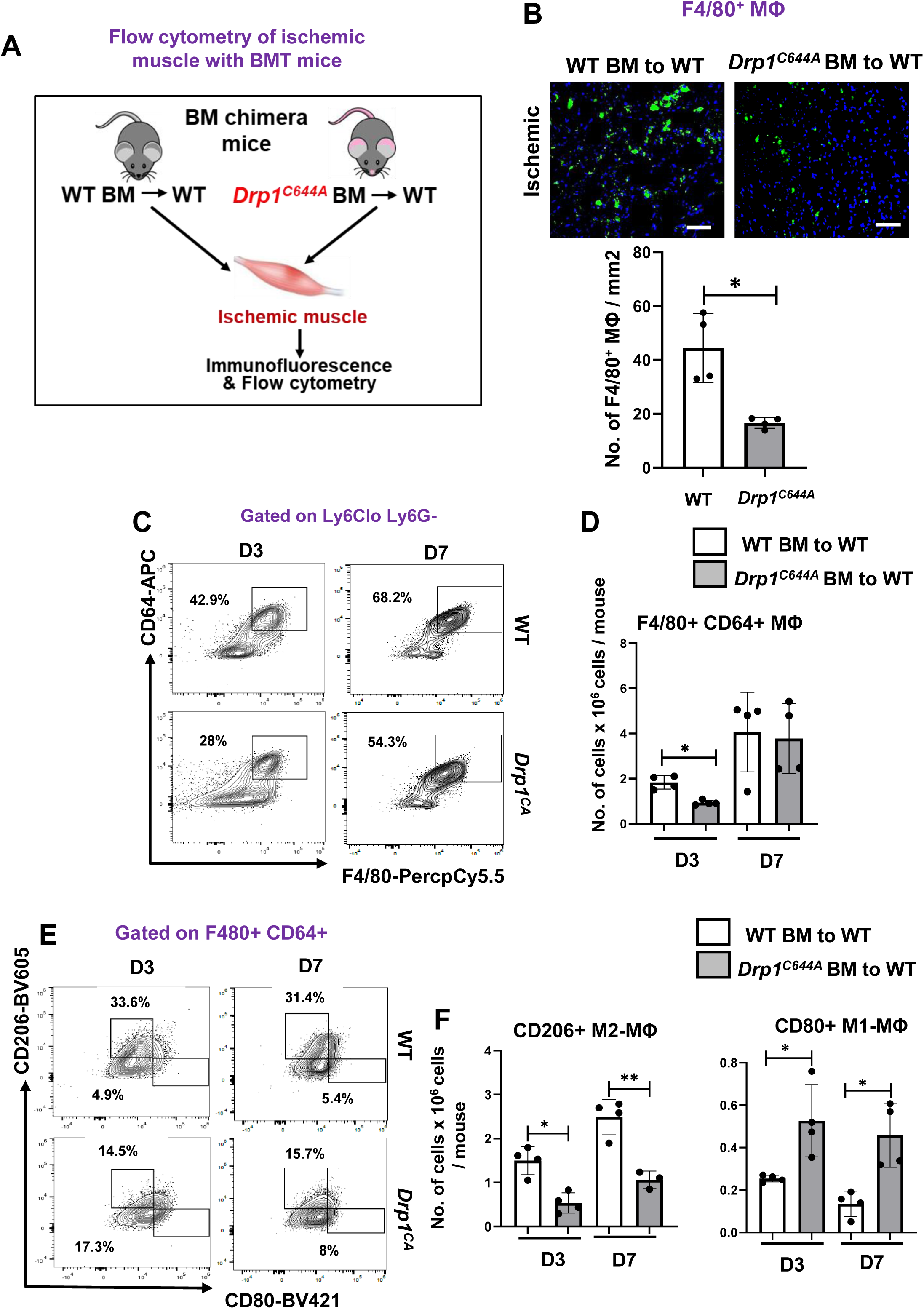
*Drp1 ^C/A^* bone marrow chimera mice show increased pro-inflammatory M1-Mɸ polarization and decreased anti-inflammatory M2-Mɸ polarization in ischemic muscles after HLI. **(A)** Schematic of flow cytometry based immunophenotyping of ischemic gastrocnemius muscle (GC) after BMT and HLI surgery. **(B)** Representative immunofluorescence images of F4/80^+^ macrophages and its quantification in GC muscle at day 3 after HLI. **(C-D)** Representative FACS contour plot of total F4/80^+^ CD64^+^ macrophages and its quantification. **(E-F)** CD80^+^M1- and CD206^+^M2-MΦ and its quantification in ischemic GC muscle at day 3 and day 7 post HLI. Data are mean ± SEM (n=4 mice). *P<0.05, **P<0.01.

### 3.3. Inhibition of ischemia-induced DRP1 sulfenylation in macrophages promotes mitochondrial hyperfusion and excess mitoROS generation *in vitro*

To define the mechanism by which DRP1 cysteine oxidation regulates macrophage polarization under ischemic conditions, we performed *in vitro* studies using BMDMs isolated from WT and *Drp1^C/A^* KI mice and exposed to hypoxia-serum starvation (HSS), an established *in vitro* model of PAD [21](Figure 3A). DCP-Bio1 assay revealed that DRP1-CysOH formation was robustly induced in WT BMDMs within 1-2 h of HSS stimulation, whereas this response was completely abolished in *Drp1^C/A^* BMDMs (Figure 3B-C). In parallel, cytosolic H₂O₂ levels were significantly increased in WT BMDMs after 1 h of HSS compared with normoxic controls (Figure S2A). Previous study shows that HSS induced a transient increase in mitochondrial fission in BMDMs, peaking at 1-2 h and returning to baseline by 8 h, without changes in canonical DRP1 phosphorylation at Ser616 or Ser637, total DRP1 expression, or levels of other mitochondrial fission and fusion proteins (MFF, MFN1/2, OPA1)[21]. In contrast, *Drp1^C/A^* KI BMDMs displayed elongated, hyperfused mitochondria at 2 h under HSS (Figure 3D-E), closely resembling the phenotype observed in Drp1-deficient BMDMs [21]. We next assessed whether loss of DRP1-CysOH alters mitochondrial function by measuring oxygen consumption rate (OCR) using Seahorse analysis. Under HSS conditions, *Drp1^C/A^*KI BMDMs showed no significant differences in basal respiration, maximal respiration, or ATP production compared with WT BMDMs (Figure S3A). Despite preserved mitochondrial respiratory capacity, *Drp1^C/A^* KI BMDMs exhibited markedly increased mitochondrial ROS production after 2 h of HSS, as detected by MitoSOX fluorescence (Figure 3F-G). Together, these results indicate that ischemia-induced DRP1 sulfenylation is required for proper mitochondrial fission in macrophages. Loss of DRP1-CysOH leads to mitochondrial hyperfusion and excessive mitoROS generation under ischemic stress, providing a mechanistic link between redox regulation of DRP1, mitochondrial dynamics, and macrophage dysfunction in PAD.

**Figure 3:**
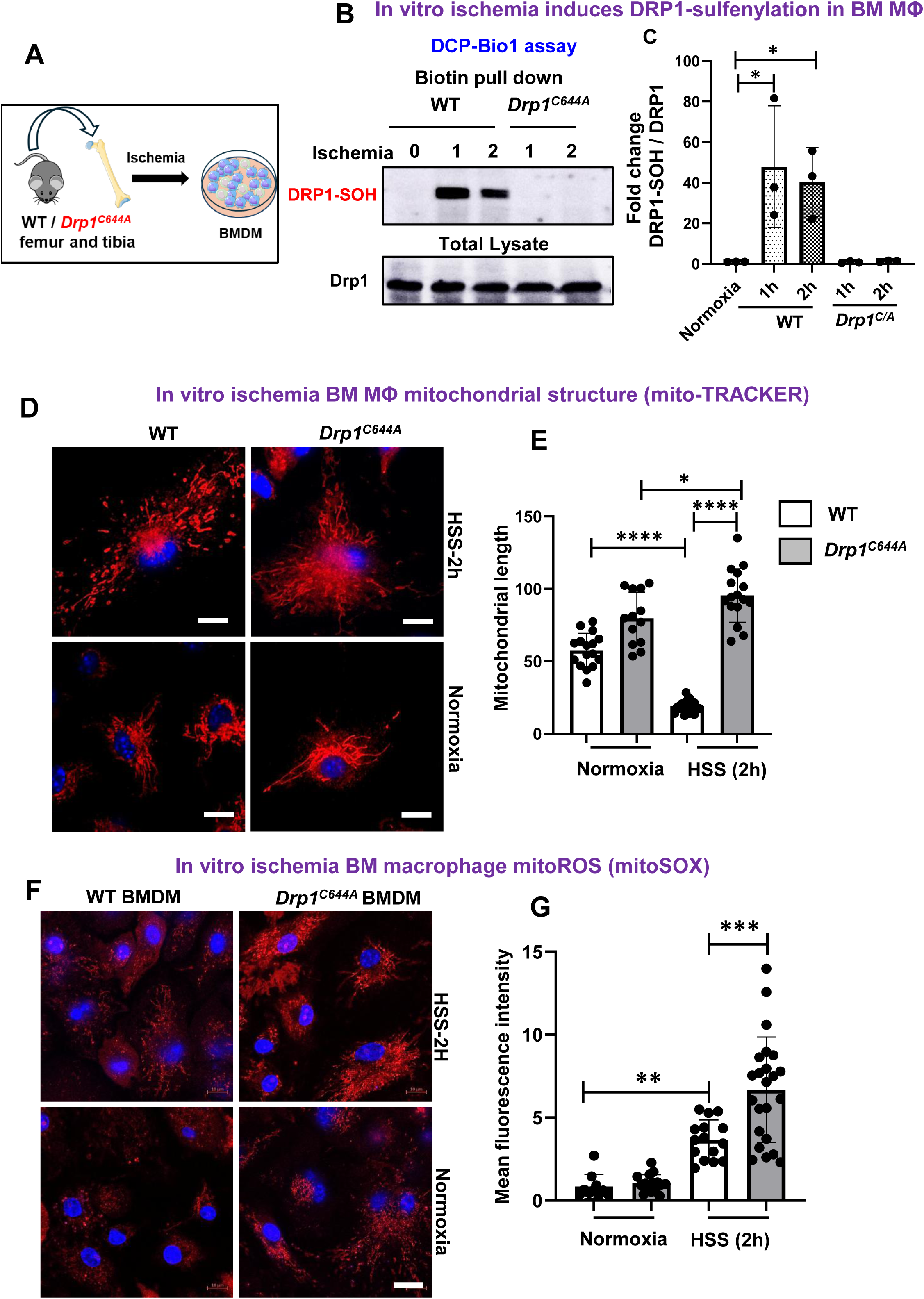
Preventing DRP11-CysOH formation in *Drp1^C/A^* KI BMDMs during ischemia induced mito-fusion and excess mitoROS generation. **(A)** Schematic of in vitro BMDM Hypoxia Serum Starvation (HSS) model. **(B-C)** Immunoblotting for DRP1sulfenylation using DCP-Bio1 in WT and *Drp1^C/A^* BMDMs at indicated time post hypoxia serum starvation (HSS). **(D-E)** mitoTracker and **(F-G)** mitoSOX staining of mitochondria and mitoROS respectively and its quantification in WT and *Drp1^C/A^* BMDM post 2h of HSS stimulation. Data are mean ± SEM (n=2). *P<0.05, **P<0.01, ***P<0.001, ****P<0.0001

### 3.4. Inhibition of ischemia-induced DRP1 sulfenylation suppress AMPK activation, enhances glycolysis and skews macrophage polarization to pro-inflammatory phenotype

To directly determine the role of DRP1-CysOH in macrophage metabolic reprogramming and polarization under ischemic conditions, BMDMs were subjected to HSS *in vitro*. Because ischemia-driven pro-inflammatory macrophage polarization is associated with a metabolic shift from oxidative phosphorylation to aerobic glycolysis [25,26], glycolytic activity was assessed by measuring extracellular acidification rate (ECAR) using a Seahorse assay. As shown in Figure 4A, *Drp1^C/A^* KI BMDMs exhibited significantly increased basal glycolysis and glycolytic capacity compared with WT BMDMs under HSS. To elucidate the mechanism underlying enhanced glycolysis in *Drp1^C/A^* KI macrophages, we examined AMP-activated protein kinase (AMPK) signaling, which promotes anti-inflammatory, pro-angiogenic M2 polarization in part by suppressing glycolysis, as previously reported in *Drp1^⁻/⁻^* BMDMs [21]. Consistent with this pathway, *Drp1^C/A^* KI BMDMs displayed markedly reduced AMPK phosphorylation and a modest increase in NF-κB phosphorylation compared with WT controls under HSS (Figure 4B). In parallel with our in vivo hindlimb ischemia findings, *Drp1^C/A^* KI BMDMs exposed to HSS showed significantly elevated expression of M1-associated genes (Nos2, Ptgs2, and Tnfα) and reduced expression of the M2 marker Retnla relative to WT BMDMs (Figure 4C). These data indicate that myeloid DRP1-CysOH deficiency promotes glycolytic reprogramming and NF-κB activation while suppressing AMPK signaling, thereby driving pro-inflammatory M1-like macrophage polarization and impairing M2-like macrophage polarization under ischemic conditions *in vitro*.

**Figure 4:**
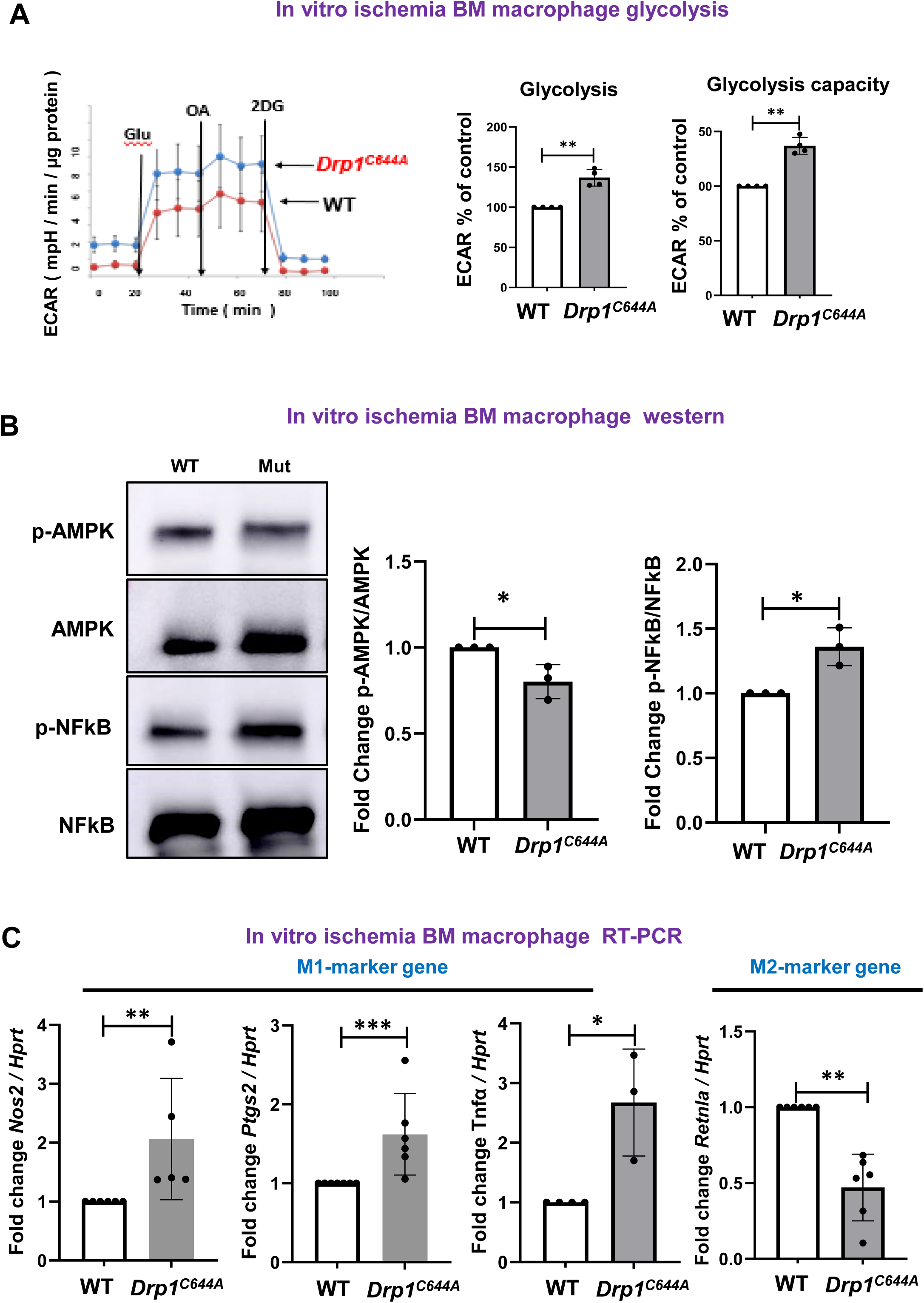
*Drp1 ^C/A^* BMDM suppressed ischemia-induced p-AMPK and increased glycolysis, pro-inflammatory M1-genes and decreased anti-inflammatory M2-gene after HSS. WT and *Drp1^C/A^* BMDM after 2 h of HSS were used to measure (**A**) glycolysis using Seahorse analyzer **(B)** p-AMPK, p-NFkB, total AMPK and NFkB using immunoblotting. **(C)** M1- and M2-marker genes normalized to *Hprt* after 8 h of HSS by qRT-PCR analysis. Data are mean ± SEM. (n=3-5). *P<0.05, **P<0.01, ***P<0.001.

**Figure 5:**
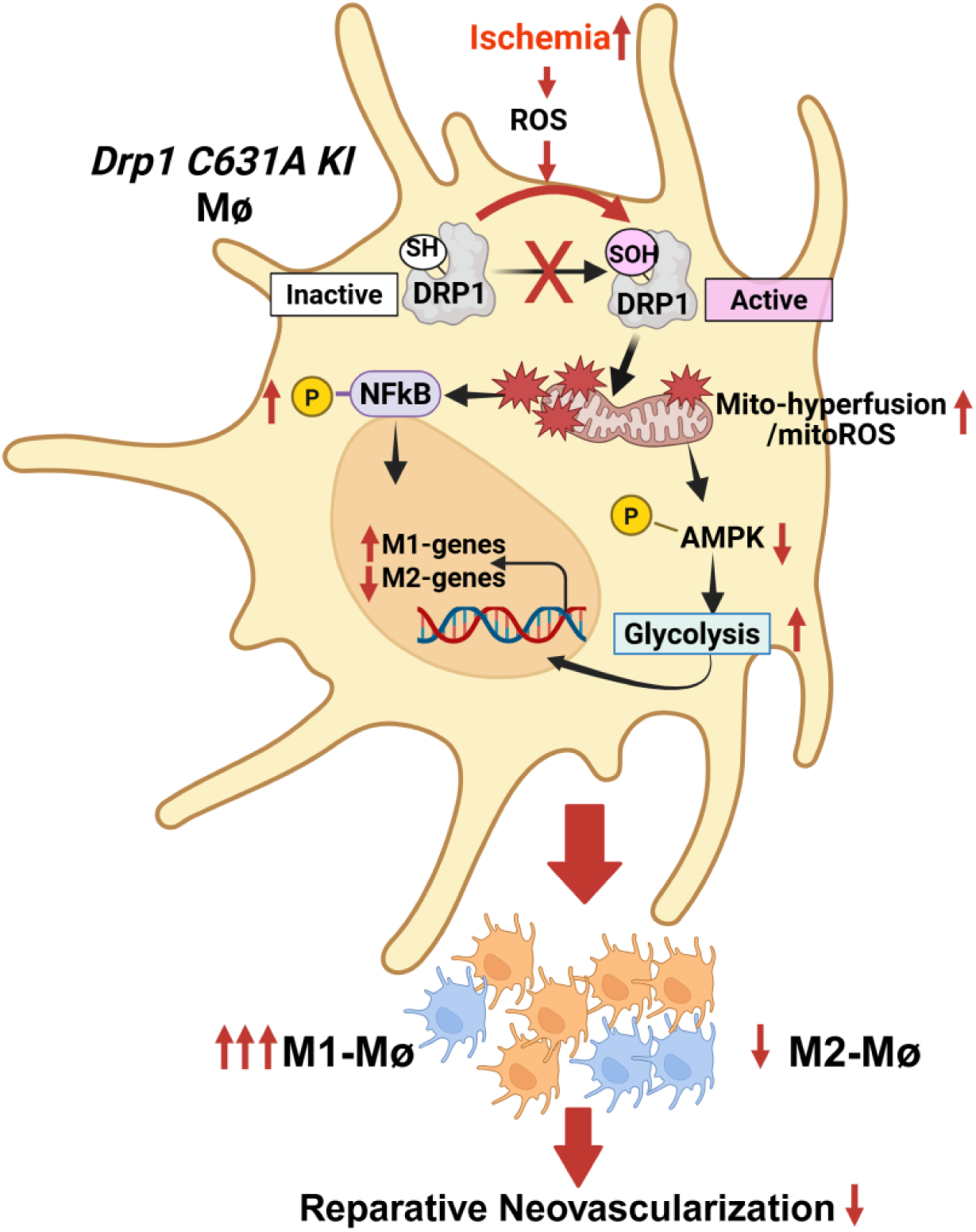
Schematic proposed model. Myeloid DRP1 sulfenylation at Cys^631^ functions as a redox sensor linking ischemia-induced ROS to drive reparative macrophage reprograming and revascularization following HLI. Loss of Drp1 sulfenylation in redox-dead DRP1^C631A^ macrophages leads to excess mitoROS, glycolysis, NFκB activation and pro-inflammatory M1 polarization while suppressing reparative AMPK activation and M2 polarization, which limits revascularization in experimental PAD.

## 4. Discussion

ROS derived from BM cells in addition to ECs are key signaling mediators required for ischemia-induced neovascularization in PAD [6,7,10,22]. We previously showed that ischemia activates the mitochondrial fission protein DRP1 in macrophages independently of canonical Ser616/Ser637 phosphorylation, but the underlying redox mechanism was unknown. Here, we identify ischemia-induced cysteine sulfenylation of DRP1 as an essential redox modification in BM cells especially macrophage promoting effective perfusion recovery and angiogenesis after HLI. BM chimeric mice expressing a “redox-dead” *Drp1^C/A^*KI mutant displayed impaired perfusion recovery and angiogenesis, accompanied by excessive neutrophil accumulation, sustained pro-inflammatory M1 macrophage polarization, and reduced M2 macrophage responses in ischemic muscle. Mechanistically, ischemic stress induced ROS-dependent DRP1 sulfenylation and mitochondrial fission without altering DRP1 phosphorylation, whereas loss of DRP1 cysteine oxidation disrupted mitochondrial dynamics, enhanced glycolysis, mitoROS and NF-κB signaling, suppressed AMPK activation, and skewed macrophages toward a pro-inflammatory state.

DRP1 is a central mediator of mitochondrial fission, and its activity is tightly controlled by multiple post-translational modifications (PTMs), including phosphorylation, SUMOylation, ubiquitination, S-nitrosylation, and O-GlcNAcylation, which collectively fine-tune mitochondrial and cellular dynamics [27]. We previously reported that ischemia activates DRP1 in macrophage without phosphorylation, which is required for reparative neovascularization [21]. In the present study, we demonstrate for the first time that DRP1-CysOH formation is dramatically increased in BM cells following HLI. Importantly, BM chimera mice harboring the redox-dead DRP1-C^644^A KI mutation exhibits impaired limb perfusion recovery and angiogenesis after HLI, establishing a functional requirement of DRP1 sulfenylation in ischemic revascularization. Furthermore, *in vitro* ischemia increased mitochondrial fission, H₂O₂ levels and DRP1-CysOH formation in BMDMs, independently of phosphorylation at Ser616 or Ser637 [21], confirming that macrophage DRP1 is activated primarily through Cys^644^ sulfenylation rather than canonical phosphorylation pathways. Although the precise ROS source driving ischemia-induced DRP1 sulfenylation in macrophages was not directly examined, our prior studies have identified NOX2 as a dominant source of ROS regulating redox signaling in BM cells during HLI [6,8,9,28]. Notably, we showed that NOX2-derived ROS from infiltrating inflammatory cells and ECs are required for neovascularization in ischemic muscle [8], and that increased NOX2-derived ROS in the BM microenvironment promotes tissue repair by regulating myelopoiesis and progenitor cell activity during post-ischemic repair [9]. Together, these findings support a model in which NOX2-derived ROS generated during ischemia are sensed by myeloid DRP1 via Cys^644^ sulfenylation, thereby promoting reparative revascularization.

Ischemia-induced revascularization depends on pro-angiogenic and metabolic reprogramming of macrophages, in which coordinated shift between oxidative metabolism and glycolysis determines inflammatory fate and reparative function [15]. Our findings identify myeloid DRP1 sulfenylation as a previously unrecognized mechanism that links redox signaling to pro-angiogenic macrophage reprogramming through control of mitochondrial dynamics, driving post-ischemic neovascularization. Consistent with our previous findings in *MΦDrp1^-/-^* mice [21], *Drp1^C/A^* KI BM chimera mice exhibited excessive neutrophil accumulation and reduced macrophage accumulation at day 3 after HLI without affecting monocyte infiltration. This was accompanied by a sustained shift toward pro-inflammatory CD80⁺ M1-like polarization and reduced CD206⁺ M2-like macrophages at both day 3 and day 7 post-HLI. Although macrophage numbers recovered by day 7, defective polarization persisted, indicating that DRP1 sulfenylation primarily regulates macrophage functional fate rather than recruitment. In addition, a rapid increase in neutrophils accumulation in *Drp1^C/A^* KI BM chimera mice after HLI suggests a role for DRP1 sulfenylation in promoting efferocytosis of apoptotic neutrophils, leading to M2 macrophage polarization in ischemic tissues [29–31]. Addressing underlying mechanisms is the subject of future studies. Mechanistically, consistent with our prior work with *Drp1^-/-^* macrophages under *in vitro* ischemic conditions [21], the present findings suggest that loss of DRP1-CysOH formation under ischemia enhanced glycolysis via suppressed AMPK activation and induced excessive mitochondrial ROS production and activation of NF-κB signaling in macrophage. This in turn drives sustained M1-like macrophage polarization and impairing revascularization in experimental PAD. These findings implicate DRP1 as a redox-sensitive mitochondrial checkpoint that integrates ROS signaling with metabolic and inflammatory pathways to determine macrophage functional outcomes during ischemic repair.

These are several limitations of the present study. First, while our data supports a central role for ROS-dependent DRP1 sulfenylation in macrophage reprogramming during ischemia, the precise upstream sources of ROS that mediate DRP1 Cys oxidation in macrophages were not directly defined. Future studies using cell-specific genetic or pharmacologic targeting of candidate ROS sources, such as NOX2 will be important to delineate the redox signals that activate DRP1 *in vivo*. Second, the molecular mechanisms linking DRP1-oxidation-dependent mitochondrial dynamics to pro-angiogenic and metabolic reprograming remain to be elucidated. Finally, whether DRP1 redox regulation similarly controls macrophage function in other ischemic or inflammatory disease contexts, including diabetic PAD or aging-associated vascular dysfunction, warrants further investigation. Addressing these questions will not only refine the mechanistic framework of DRP1 redox signaling but also inform the translational potential of targeting DRP1 sulfenylation to enhance therapeutic revascularization in PAD.

## Conclusions

In conclusion, our study uncovers a previously unrecognized mechanism by which ischemia-derived ROS are sensed to promote reparative angiogenic responses through DRP1 cysteine sulfenylation in macrophages. By uncoupling DRP1 activation from canonical phosphorylation and identifying sulfenylation as the dominant regulatory mode in ischemic macrophages, these findings redefine how macrophage mitochondrial dynamics are controlled during tissue repair. Targeting DRP1 redox signaling may therefore represent a novel therapeutic strategy to enhance macrophage-mediated revascularization while limiting chronic inflammation in PAD.

## Author Contributions

M.U.-F., T.F. and S.Y. designed the study; S.Y, S.N., S.V., R.K., S.K., A.D., performed/assisted research; M.U.-F., T.F., S.Y. analyzed data; V.G., S.K. performed mouse genotyping; M.U.-F., T.F. and S.Y. wrote the manuscript.

## Acknowledgements

We acknowledge Dr. Lin Gan at Transgenic & Genomic Editing Core at Augusta University Medical College of Georgia for generating Drp1-C^631^A KI mutant mice.

## Data Availability Statement

The data presented in this study are available in the article and supplementary materials.

## Supplementary Figure legends

**Figure S1.**
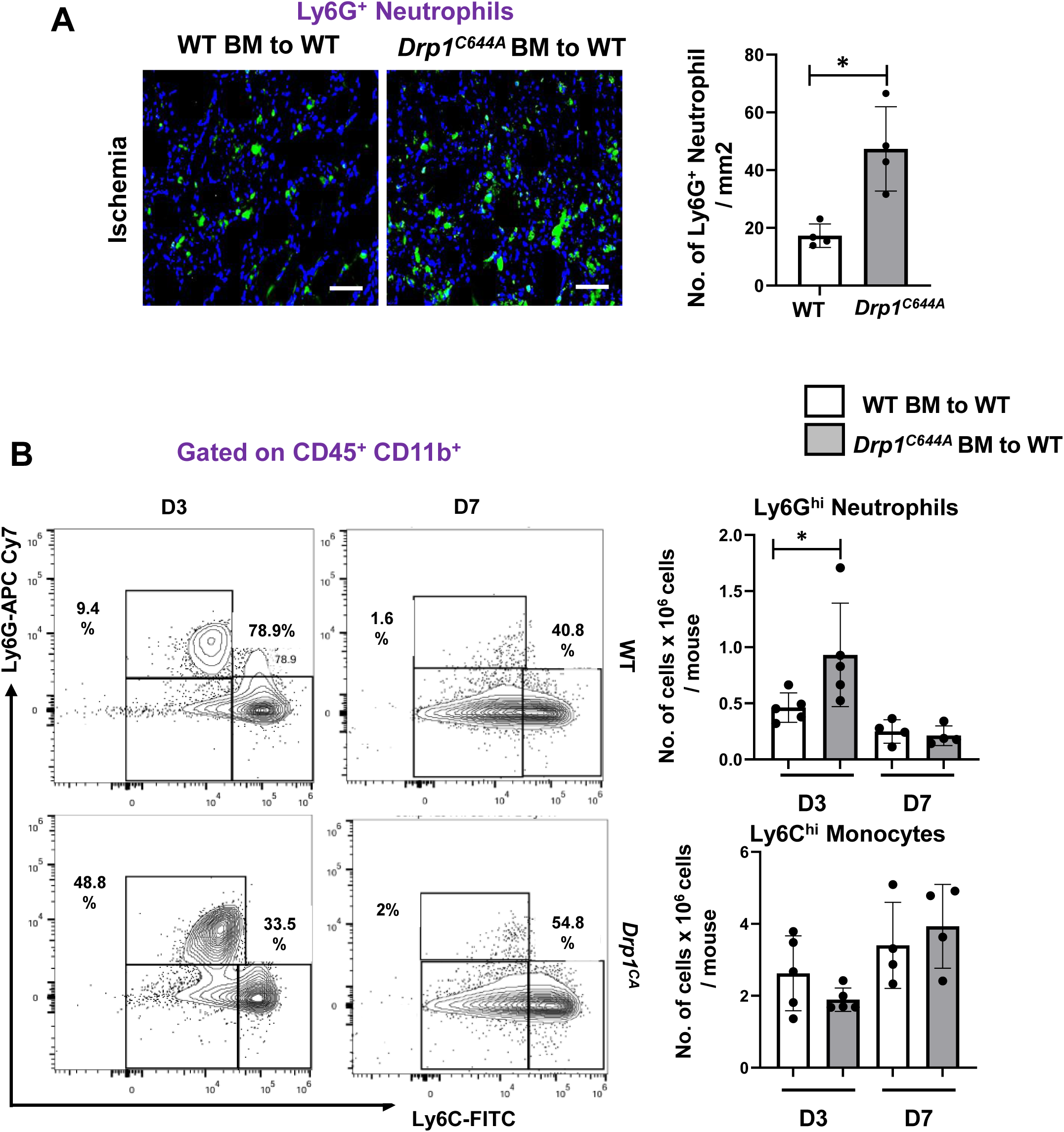
Neutrophils and monocyte recruitment in ischemic muscles of WT and *Drp1^C/A^* BM chimera mice after HLI. **(A)** Representative immunofluorescence images of Ly6G^+^ neutrophils and its quantification in GC muscle at day 3 after HLI. **(B-C)** Representative flow cytometry contour plots and quantification of numbers of Ly6G^hi^ neutrophils and Ly6C^hi^ monocytes in ischemic GC muscle of WT and *Drp1^C/A^*mice after day 3 of HLI. Data are mean ± SEM. n=3-5. *p<0.05.

**Figure S2.**
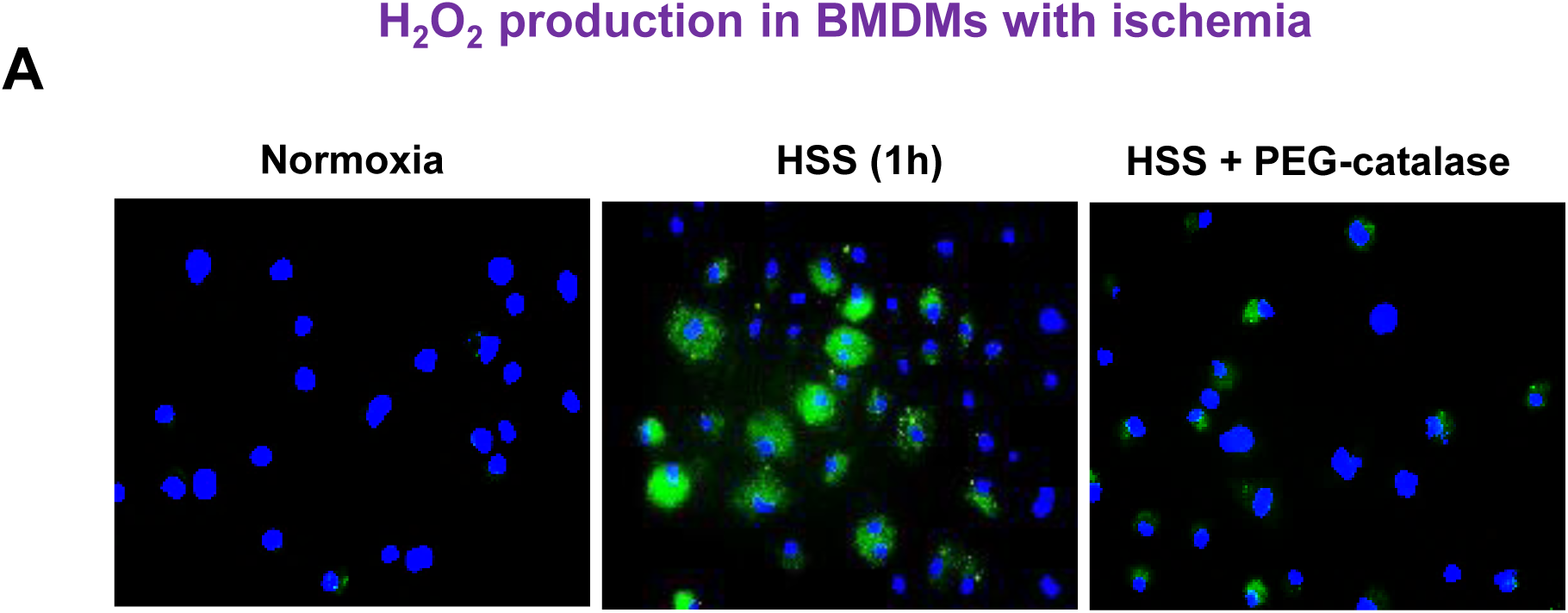
A. *In vitro* ischemia induced H_2_O_2_ in WT BMDM. DCF-DA (H_2_O_2_production) fluorescence in WT BMDMs during normoxia vs 1h of HSS stimulation with or without PEG-catalase.

**Fig. S3.**
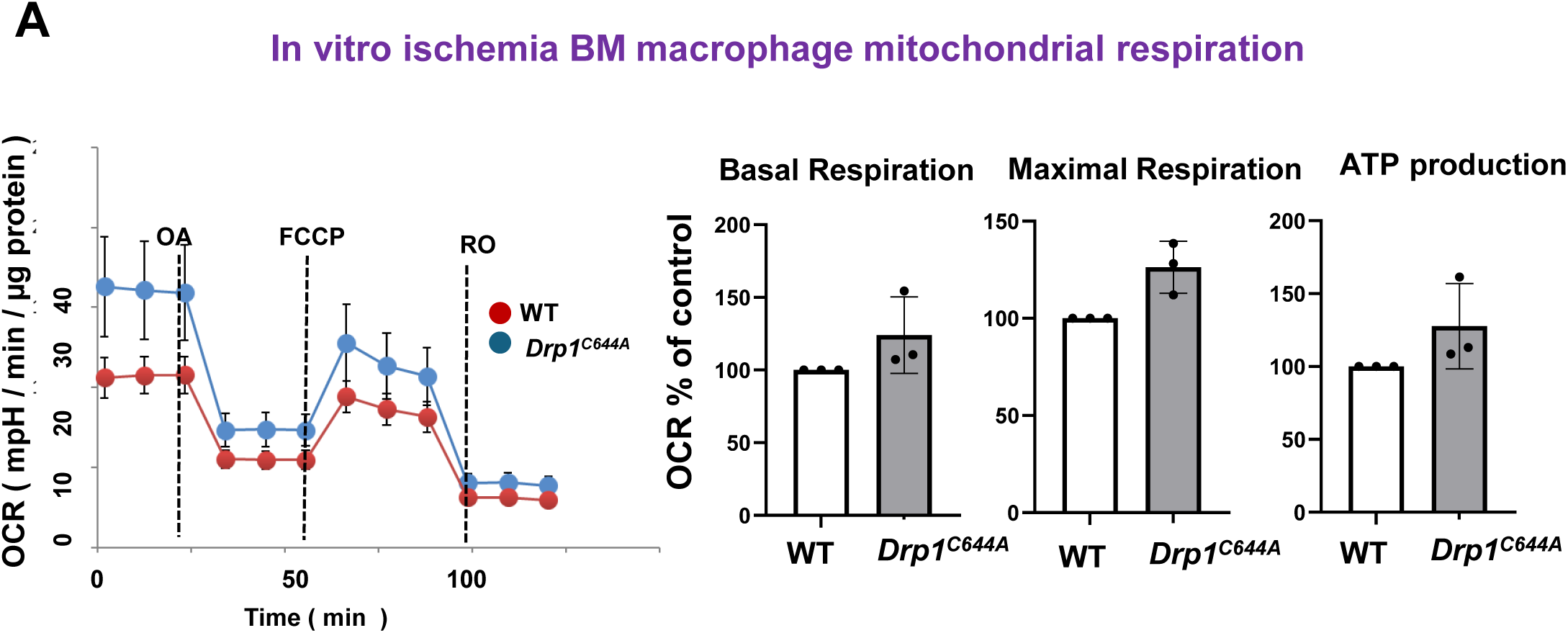
Mitochondrial respiration in *Drp1^C/A^* BMDMs with ischemia is unaffected. **A.** WT and *Drp1^C/A^* BMDMs after 2 h of HSS were used to measure mitochondrial respiration rate (OCR) using Seahorse. Data are mean ± SEM. n=2.

